# High-order epistasis in catalytic power of dihydrofolate reductase gives rise to a rugged fitness landscape in the presence of trimethoprim selection

**DOI:** 10.1101/398065

**Authors:** Yusuf Talha Tamer, Ilona K. Gaszek, Haleh Abdizadeh, Tugce Altinusak Batur, Kimberly Reynolds, Ali Rana Atilgan, Canan Atilgan, Erdal Toprak

## Abstract

Evolutionary fitness landscapes of certain antibiotic target enzymes have been comprehensively mapped showing strong high order epistasis between mutations, but understanding these effects at the biochemical and molecular levels remained open. Here, we carried out an extensive experimental and computational study to quantitatively understand the evolutionary dynamics of *Escherichia coli* dihydrofolate reductase (DHFR) enzyme in the presence of trimethoprim induced selection. Biochemical and structural characterization of resistance-conferring mutations targeting a total of ten residues spanning the substrate binding pocket of DHFR revealed distinct resistance mechanisms. Next, we experimentally measured biochemical parameters (*K*_*m*_, *K*_*i*_, and *k*_*cat*_) for a mutant library carrying all possible combinations of six resistance-conferring DHFR mutations and quantified epistatic interactions between them. We found that the epistasis between DHFR mutations is high-order for catalytic power of DHFR (*k*_*cat*_ and *K*_*m*_), but less prevalent for trimethoprim affinity (*K*_*i*_). Taken together our data provide a concrete illustration of how epistatic coupling at the level of biochemical parameters can give rise to complex fitness landscapes, and suggest new strategies for developing mutant specific inhibitors.

## Introduction

Antibiotic resistance is one of the most important global health threats [1]. According to the Centers for Disease Control and Prevention, antibiotic resistant pathogens cause over 20,000 deaths and two million infections annually in the United States alone [2]. Antibiotic resistance evolves either by resistance-conferring spontaneous mutations in bacterial genomes or horizontal transfer of mobile resistance elements [3, 4]. These genetic changes typically confer resistance by reducing the affinities of antibiotic molecules to their targets, deactivating antibiotics by chemical modification, and finally decreasing effective antibiotic concentrations inside bacterial cytoplasm by either efflux pumps or reduced uptake of antibiotic molecules [5]. Among these, understanding how mutations render antibiotics ineffective by altering their targets is particularly important from both clinical and basic science perspectives [6, 7].

In pathogenic bacteria, there is only a handful of drug target enzymes, such as DNA gyrases and RNA polymerases and finding new “druggable” enzymes or novel drugs that can target resistant bacteria is often a long and extremely difficult process [8-12]. Therefore, a mechanistic understanding of resistance-conferring mutations in already known antibiotic target enzymes is critical for designing new drugs or drug variants that can inhibit antibiotic resistant bacteria [13, 14]. How essential enzymes can preserve their catalytic activities when they acquire mutations to reduce drug affinity is another important question for better understanding basic principles driving protein evolution [7, 15-18]. In this study, we scrutinize molecular mechanisms of resistance conferring mutations in the *Escherichia coli* dihydrofolate reductase (DHFR) enzyme and investigate how epistasis between these mutations shape the adaptive landscape for trimethoprim resistance evolution.

DHFR is a ubiquitous enzyme in nature with an essential role in folic acid synthesis [19-21]. Due to its central role in metabolism (Figure 1A), DHFR is used as a drug target in antibacterial, anticancer, antirheumatic, and antimalarial therapies [21]. For instance, pyrimethamine is one of the few available drugs that can be used for treating malaria caused by *Plasmodium falciparum*, the most common species that causes malaria in humans. Pyrimethamine has specific toxicity against *P. falciparum* by binding and inhibiting the *P. falciparum* dihydrofolate reductase (pfDHFR) enzyme [13, 22, 23]. However, although pyrimethamine was one of the most commonly used drugs for malaria treatment in the past, as of today, it is rarely prescribed due to the resistance problem [22, 24]. The most common resistance-conferring mutations in pfDHFR are the four point mutations N51I, C59R, S108N, and I164L [22, 23]. The quadruple mutant of pfDHFR that carries all four of these mutations is widespread globally and is highly resistant to pyrimethamine. Similarly, evolution of resistance to trimethoprim (TMP), a bacteriostatic antibiotic molecule that competitively binds to DHFR and blocks its enzymatic activity, proceeds through sequential accumulation of resistance-conferring mutations in the bacterial DHFR enzyme [25, 26]. In our previous work, we showed that *E. coli* cells evolved trimethoprim resistance by accumulating up to four DHFR mutations in a stepwise fashion [15, 25, 26]. Since DHFR is an essential enzyme, the evolution of resistance against DHFR inhibiting drugs is a search for finding DHFR mutants that have reduced drug affinity and yet adequate catalytic power for organismal survival. For better understanding the evolutionary dynamics of resistance against DHFR inhibitors, it is important to quantitatively evaluate evolutionary paths leading to antibiotic resistance and characterize resistance at the molecular level for the ultimate goal of improving human health.

**Figure 1:**
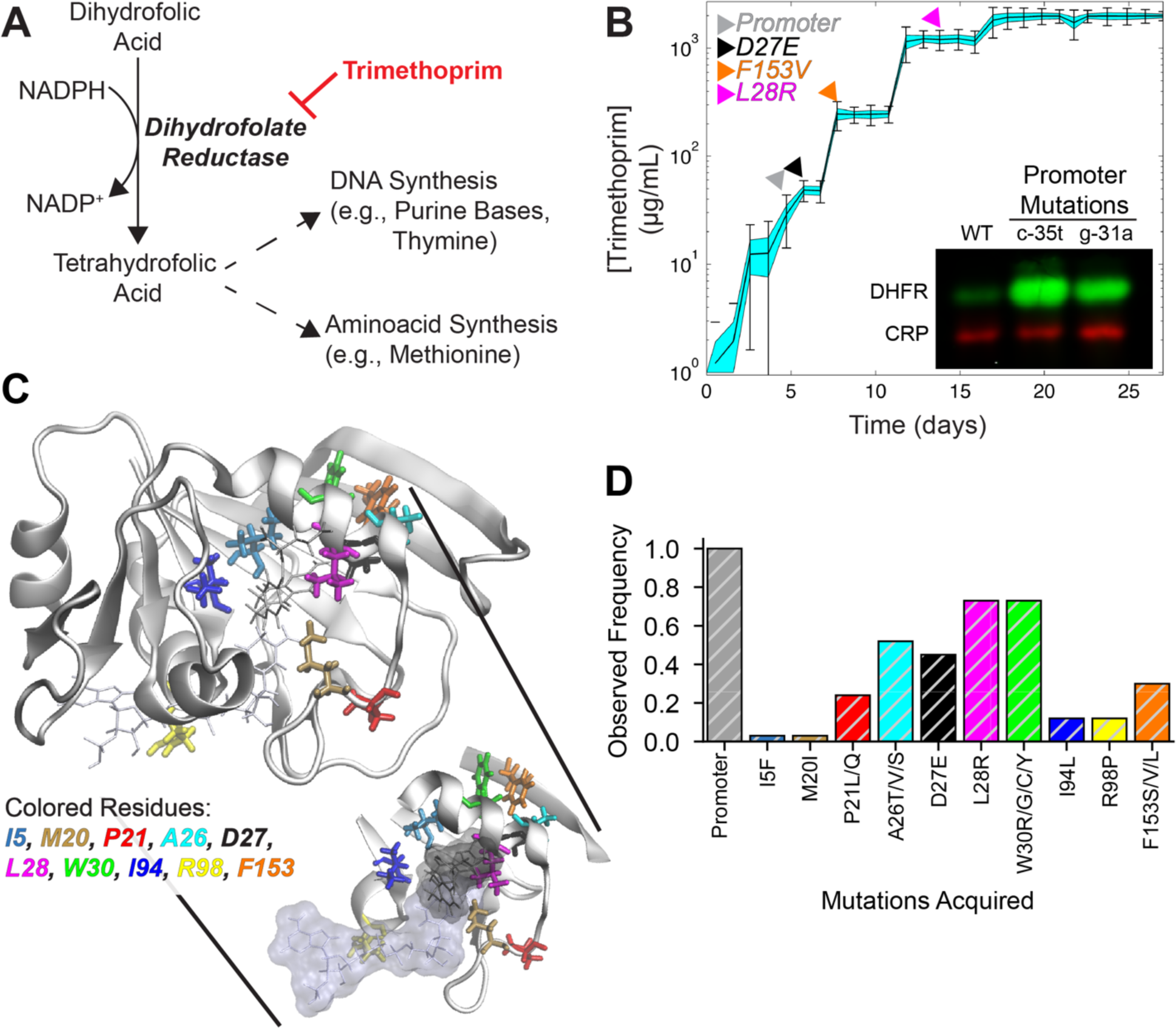
Trimethoprim resistance evolves through sequential accumulation of DHFR mutations. **A)** Enzymatic activity of DHFR is crucial for nucleotide and amino acid synthesis in *E. coli*. Trimethoprim is a competitive inhibitor of DHFR that blocks its enzymatic activity by occupying its active site. **B)** Morbidostat experiments revealed stepwise acquisition of resistance conferring mutations; a sample morbidostat trajectory demonstrating temporal changes in trimethoprim resistance. Colored arrows indicate the timing of the first detection of DHFR mutations. (Insert) Promoter mutations (c-35t, g-31a) lead to 10 to 20-fold higher DHFR expression relative to WT. **C)** Mutated DHFR residues are highlighted in different colors on DHFR structure (PDB ID: 1rx2). **D)** Observed frequencies of resistance conferring mutations plotted for 33 independent morbidostat experiments (28 populations from this study and 5 populations from a previous study [26]).

We carried out a comprehensive experimental and computational study to better understand the evolutionary dynamics of *Escherichia coli* DHFR in the presence of trimethoprim. In the following part of this text, DHFR will be used to refer *Escherichia coli* dihydrofolate reductase enzyme. We evolved several antibiotic naïve *E. coli* populations against trimethoprim in the morbidostat, a continuous culture device we developed to quantitatively study antibiotic resistance evolution [26, 27]. We identified genetic changes in *E. coli* that were responsible for trimethoprim resistance by using both whole genome sequencing and targeted gene sequencing. The genetic changes we found were almost exclusively targeting the *folA* gene that encodes for DHFR. We identified ten residues that were frequently mutated in the DHFR as well as promoter mutations that significantly increased DHFR expression. We characterized these mutations by quantifying their effects on substrate binding (*K*_*m*_), inhibitor binding (*K*_*i*_), and catalytic rate (*k*_*cat*_) of DHFR. We synthesized all possible combinations for six of these DHFR mutations and quantified epistatic interactions between these mutations. Finally, we measured the effects of these mutations on bacterial fitness by replacing the endogenous *folA* gene in *E. coli* with its mutated variants. Our analysis shows that the adaptive landscape of DHFR deviates from the landscape predicted from the fitness effects of single mutations on the wild-type DHFR using Bliss independence model where fitness effects of multiple mutations are additive. This difference is mainly because of the high-order epistasis between mutations altering DHFR catalytic activity and substrate binding. Next, we carried out molecular dynamics (MD) simulations to reveal structural changes responsible for trimethoprim resistance and epistatic interactions between mutations. Analysis of the MD simulations suggest that DHFR mutations confer resistance by utilizing distinct mechanisms which may be exploited for drug design purposes. They also point to possible dynamical mechanisms leading to epistasis. Finally, by running computer simulations, we identified plausible genetic trajectories that reach to trimethoprim resistant genotypes. Our simulations suggest that the evolution of trimethoprim resistance can be impeded by exploiting epistatic interactions between resistance-conferring mutations and the use of mutant specific inhibitors.

## Results

DHFR catalyzes the reduction of 7,8-dihydrofolate (DHF) to 5,6,7,8-tetrahydrofolate (THF) by hydride transfer from nicotinamide adenine dinucleotide phosphate (NADPH) (Figure 1A) [20, 21, 28-31]. THF is an essential precursor for cell growth as it is used in thymidylate and purine synthesis. Therefore, inhibition of bacterial DHFR slows down or stops bacterial growth. Trimethoprim is a bacterial DHFR inhibitor which competitively binds to the active site of DHFR. It is a commonly used antibiotic compound for treating bacterial infections and is typically used in combination with sulfamethoxazole due to synergism in their combined effects. We and others have previously run laboratory evolution experiments to explore evolutionary trajectories that lead to high levels of trimethoprim resistance in *E. coli* [25, 26, 32]. In these studies, we have shown that trimethoprim resistance evolved in a stepwise fashion and all populations acquired multiple mutations in the *folA* gene that encodes DHFR. One of these mutations was always in the promoter region and the rest were in the coding region of *folA*. Mutations elsewhere in the genome were rare implying that the evolution of trimethoprim resistance was confined to a small genetic target [26]. Although our results suggested a reproducibility in the temporal order of the DHFR mutations, the number of evolved populations was small and it was not clear whether the mutations we observed were covering all possible DHFR mutations. This observation was consistent with previous studies reporting multiple DHFR mutations in clinically isolated trimethoprim resistant pathogens [33, 34]. Besides, since a decrease in DHFR’s catalytic efficiency is expected to decrease bacterial fitness [35], it was not clear whether evolutionary trajectories would have been different if the minimum allowed growth rate in an evolution experiment was changed.

### Mutational trajectories observed in the morbidostat are independent of the imposed growth rate constraints

We evolved 28 initially isogenic and trimethoprim sensitive *E. coli* populations in the morbidostat using different minimum growth rate constraints [26, 27]. Morbidostat is an automated continuous culture device that maintains a nearly constant selection pressure even when bacterial populations evolve higher antibiotic resistance. This is achieved by continuously monitoring bacterial growth and clamping bacterial growth rate by adjusting antibiotic concentrations with the help of computer controlled pumps. Addition of plain growth media or antibiotic containing growth media is periodically done at constant dilution rates. Therefore, populations or subpopulations that cannot grow faster than the dilution rate of the morbidostat are washed out and hence cannot survive in the morbidostat. This feature enabled us run evolution experiments at different dilution settings and control the minimal growth rate allowed for the survival of bacterial populations. In our settings, the drug-free growth rate of the parental *E. coli* strain (MG1655) was ∼0.8 hour^−1^ (M9 minimal media supplemented with casamino acids and glucose, at ∼30°C). We evolved initially isogenic and antibiotic naïve *E. coli* populations (MG1655, Materials and Methods) at three different dilution rates (0.3 h^−1^ (n=7), 0.45 h^−1^ (n=7), 0.6 h^−1^ (n=14)) for several weeks and asked whether there would be any difference in the evolutionary dynamics of trimethoprim resistance.

All *E. coli* populations evolved very high trimethoprim resistance in a stepwise fashion (Figure 1B) and they were able to survive even at ∼3 mg/ml trimethoprim concentration which is the maximum solubility limit of trimethoprim in our growth media (M9 minimal media supplemented with casamino acids and glucose, at 30°C). All of the populations acquired three to five mutations in the *folA* gene and whole genome sequencing of 15 randomly selected mutants that were isolated on the last day of morbidostat experiments revealed few mutations elsewhere in the genome (Table S1). One of the mutations in the *folA* was always a promoter mutation (g-9a, c-15a, g-31a, c-35t) and these promoter mutations were increasing DHFR levels 10-20 times compared to their wild type ancestor (Figure 1B, insert). The rest of the folA mutations were in the coding region of *folA* and targeted total of ten residues that were spanning the substrate binding pocket as illustrated in Figure 1C. Among these, the most common mutations were at the following residues: P21, A26, D27, L28, W30, and F153 (Figure 1D). However, contrary to our expectations, we did not observe any evolutionary pattern indicating that mutations or mutational trajectories were specific to the growth rate constraints we imposed by varying dilution rates in the morbidostat. This observation suggested that the acquired DHFR mutations did not have significant effects on the bacterial growth or DHFR mutations that could diminish bacterial growth never reached to detectable levels throughout morbidostat experiments. An alternative explanation could be that the DHFR enzyme already has the capacity to tolerate catalytic deficiencies due to resistance-conferring mutations. This can be either because the DHFR already produces more THF than required for growth or these deficiencies were compensated by the overexpression of DHFR or by the emergence of other mutations. We conclude that *E. coli* populations evolving in the morbidostat can acquire three to five mutations to render trimethoprim ineffective and there were no patterns in the evolutionary trajectories specific to the growth rate constraints we imposed throughout the experiments.

### Resistance-conferring mutations have diverse effects on catalytic efficiency of DHFR

Ideally, fitness effects of mutations should be measured at the organismal level. However, characterizing the evolutionary fitness landscape for DHFR requires reliable fitness measurements which is not always possible when *in vivo* assays are utilized. First, in our experience, several of the bacterial mutants carrying DHFR mutations survived even at the highest possible trimethoprim concentrations we could achieve (∼3mg/ml) making it impossible to measure their true resistance levels [15]. Second, despite our numerous attempts, it was not possible to engineer some of the *E. coli* strains with desired combinations of DHFR mutations, implying that cells with some *DHFR* alleles may not be viable [15]. Third, the strain we engineered by replacing the endogenous *folA* (the gene that is transcribed into DHFR) with the wild-type *folA* gene had a growth defect compared to its ancestor MG1655 strain making growth rate measurements less reliable. Fourth, overexpression of DHFR due to promoter mutations masked the true fitness effects of mutations found in the coding region of *DHFR* [15]. Finally, it is difficult to unequivocally attribute the effects of mutations to bacterial fitness as bacterial cells can compensate deleterious effects of DHFR mutations by gene regulation or rearranging metabolic fluxes. Therefore, we decided to characterize fitness effects of DHFR mutations at the protein level by utilizing *in vitro* assays.

We developed a rapid assay for calculating *k*_*cat*_, *K*_*m*_, and *K*_*i*_ values for mutant DHFR enzymes (Figure 2). Measuring substrate affinity (*K*_*m*_) and catalytic rate (*k*_*cat*_) of an enzyme typically requires enzymatic activity measurements at various substrate concentrations and predicting *k*_*cat*_ and *K*_*m*_ values by fitting a Michelis-Menten function to the resulting data [7, 35, 36]. Depending on the enzyme, this can be a laborious and expensive task. In the case of DHFR, the standard assay used for measuring DHFR activity benefits from spectroscopic changes in the cofactor (NADPH) and substrate (DHF) of DHFR as THF is produced. Typically, by maintaining a high concentration of NADPH compared to the DHF, initial reduction rate of DHFR is calculated by monitoring the absorbance of NADPH and DHF at 340 nm wavelength. NADPH and DHF have high absorptions at 340nm (A_340_) but their absorptions become insignificant upon hydride transfer between them. When DHFR is mixed with NADPH and DHF, A_340_ rapidly reduces until DHF is completely consumed and this measurement needs to be repeated at several different substrate concentrations for predicting *k*_*cat*_ and *K*_*m*_ values. We realized that this laborious assay was not necessary for characterizing DHFR. In the presence of saturating concentrations of DHF (10-20μM) and NADPH (100-200μM), DHFR molecules already sample all possible concentrations of DHF throughout the progression of the reaction while NADPH levels are still at saturating levels.

**Figure 2:**
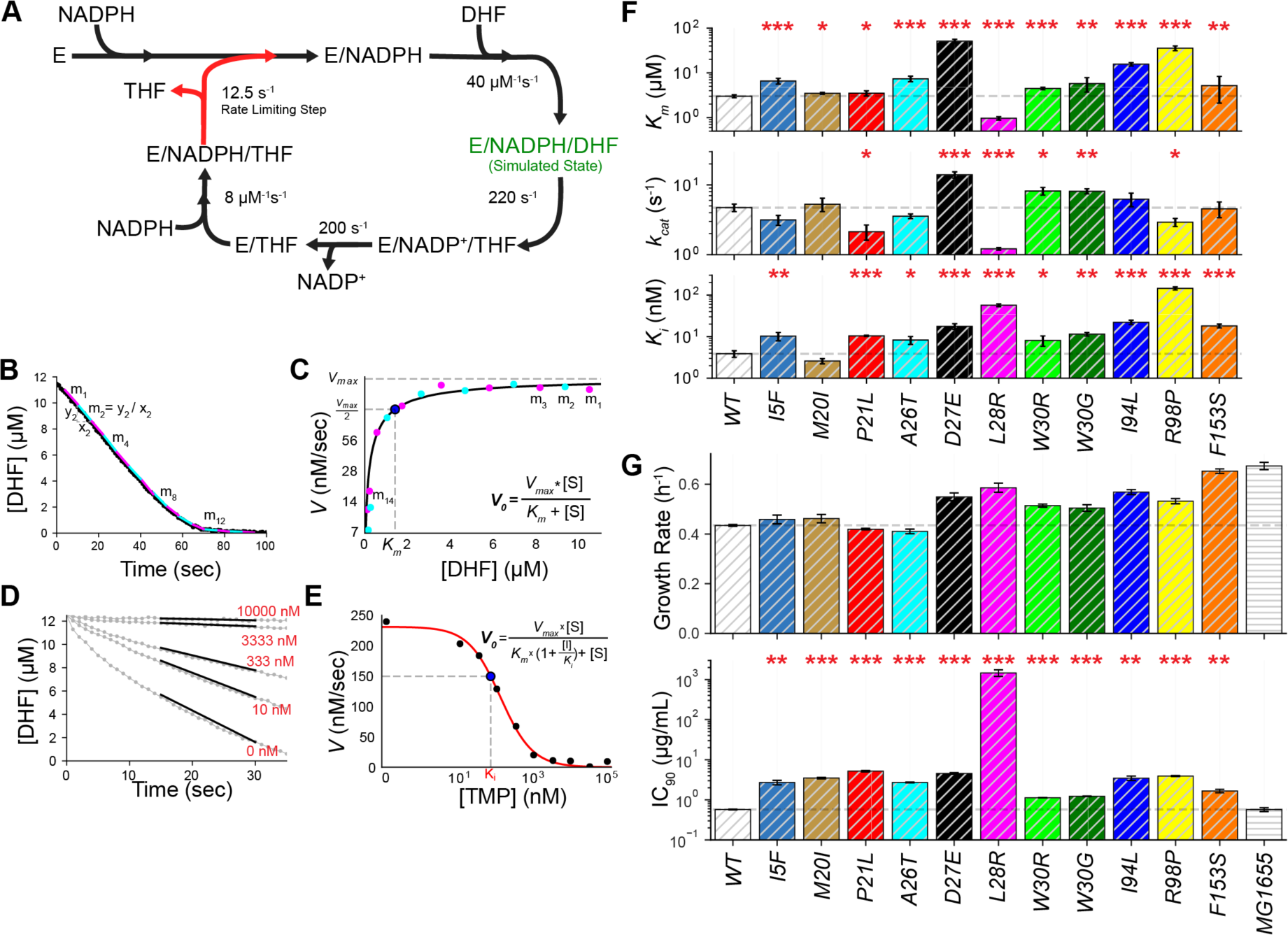
Biochemical characterization of resistance-conferring DHFR mutations. **A)** Catalytic cycle of DHFR. Forward reaction rates are obtained from Schnell et al. [21]. Rate limiting step in the catalytic cycle is release of THF (red arrow). E stands for DHFR. E-NADPH-DHF (green fonts) is the state used in our molecular dynamics simulations. **B)** Left panel shows a typical reaction progression curve after absorbance (340 nm) values are converted to DHF concentration (see Methods). By utilizing moving time windows, we calculate catalysis rates at corresponding DHF concentrations. **C)** *K*_*m*_ and *k*_*cat*_ values are predicted by fitting a Michelis-Menten equation to measured catalysis rates. **D-E)** Initial reaction rates in the presence of various trimethoprim concentrations are used to predict the affinity (*K*_*i*_) of DHFR mutants to trimethoprim molecules. **F)** *K*_*m*_, *k*_*cat*_ and *K*_*i*_ values of DHFR mutants with single amino acid replacements. Error bars show standard error of the mean. Student’s t-test (two tailed) is used to quantify significance of *K*_*m*_, *k*_*cat*_ and *K*_*i*_ changes relative to the wild type (WT) DHFR (*: p<0.05; **: p<0.01; ***: p<0.001). **G) (Upper Panel)** All engineered *E. coli* strains carrying single DHFR mutations are viable. Endogenous *folA* gene was replaced with the wild-type (WT) or mutated *folA* genes (Materials and Methods). Cells were grown at ∼30^°^C in minimal M9 media supplemented with 0.4% glucose and 0.2% amicase in 12 replicates. Exponential growth rates of all mutants except the I5F and L28R are all significantly lower than the parental MG1655 *E. coli* strain but higher that the strain (WT) we engineered by reinserting the wild-type (WT) folA gene. Despite our several attempts, the engineered WT strain had a growth defect most likely as a result of the selection markers we used for cloning (Materials and Methods). (Lower Panel) All engineered *E. coli* strains carrying single DHFR mutations have elevated trimethoprim resistance. Inhibitory concentrations reducing growth by ninety percent (IC_90_) were measured by growing mutants in a gradient of trimethoprim using 12 replicates (∼30^°^C in minimal M9 media supplemented with 0.4% glucose and 0.2% amicase). Student’s t-test (two tailed) is used to quantify significance of IC_90_ changes relative to the wild type (WT) DHFR (*: p<0.05; **: p<0.01; ***: p<0.001, error bars shows the standard error on the mean for each mutant).

Also, the spectroscopic properties of NADPH and DHF allow us to predict both DHF and NADPH concentrations during the progression of this reaction. Since the rates of reverse reactions (Figure 2A, counterclockwise direction) in the catalytic cycle are very slow relative to the forward reaction rates (Figure 2A, clockwise direction), it is possible to calculate reaction rates at various DHF concentrations from a single reaction progression curve. As shown in Figure 2B, we split the progression curve in equal time windows and calculate corresponding mean DHF concentrations and DHF reduction rates for every time interval. We then use these values to predict *k*_*cat*_ and *K*_*m*_ values by fitting a Michelis-Menten equation (Figure 2C). The *K*_*m*_ values we measured using this practical method were consistent with the values we measured using the standard conventional method that needs measurements at several different DHF concentrations (*K*_*m*_ predicted using traditional method: 3.40±0.95μM, and our method gives: 2.86±0.97μM). In addition, by measuring DHFR activity at steady state using various trimethoprim concentrations (Figure 2D), we calculated trimethoprim (TMP) affinities of DHFR mutants (*K*_*i*_) assuming competitive binding kinetics between DHF and TMP (Figure 2E, equation 1).

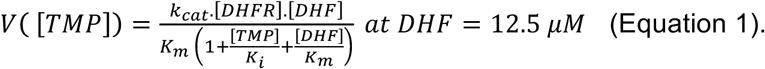

All of the mutations except the L28R caused significant reductions in the substrate affinity (increased *K*_*m*_) of DHFR (Figure 2F, Table S2). Contrary to our expectations, substrate affinity of the L28R mutant was significantly increased (decreased *K*_*m*_) relative to the wild type DHFR. Changes in the *K*_*m*_ were generally accompanied with significant changes in the *k*_*cat*_ values. Interestingly, three of the mutants (P21L, L28R, and R98P) exhibited decreased catalytic rates whereas others (D27E, W30G, and W30R) had increased catalytic rates *k*_*cat*_. Finally, all of the mutations but one (M20I) had reduced trimethoprim affinity (increased *K*_*i*_). Although antibiotic resistance via target modifications is typically attributed to reduced drug and substrate affinities due to mutations, our measurements summarized in Figure 2F suggest that there could be distinct resistance mechanisms. That being said, *K*_*i*_ values alone are far from enough for explaining trimethoprim resistance [7]. In the bacterial cell, several other parameters such as expression of DHFR, catalytic efficiency (*k*_*cat*_/*K*_*m*_), thermal stability, availability of nutrients and metabolites, accumulation of excess DHF, and the need for THF can contribute to bacterial fitness in the presence of trimethoprim. Finally, we engineered mutant *E. coli* strains by replacing wild-type *folA* gene with its variants with single mutations. All of the engineered E. coli strains with single DHFR mutations were viable (Figure 2G) and had elevated trimethoprim resistance compared to their parental MG1655 strain (Figure 2H).

In summary, all DHFR mutations except the L28R and M20I mutations decreased both substrate and inhibitor binding with the exception of M20I which did not have a significantly different *K*_*i*_ value compared to the wild-type DHFR. On the other hand, the L28R mutation increased substrate affinity and decreased catalytic rate suggesting the existence of newly formed interactions between the mutated DHFR protein and its substrate (DHF). The catalytic rates of other DHFR mutants exhibited both decreasing and increasing phenotypes. We conclude that the resistance-conferring mutations in DHFR are phenotypically diverse suggesting the presence of distinct resistance mechanisms.

### Structural evaluation of DHFR with single mutations reveal distinct resistance mechanisms at the molecular level

We utilized molecular dynamics (MD) simulations in order to study the structural changes associated with the trimethoprim resistance conferring mutations in DHFR resulting from point mutations discussed in the previous subsection (Figure 2F). *E. coli* DHFR is formed of eight stranded β-sheets and four contiguous α-helices [37-39]. The enzyme is divided by the active site cleft into two subdomains: the adenosine binding subdomain and the major subdomain. The former (residues 38–88) provides the binding site for the adenosine moiety of the cofactor (NADPH) and includes the CD loop (residues 64-71). The latter subdomain consists of ∼100 residues and contains three loops on the ligand binding face that surrounds the active site. These loops are known as M20 (residues 9–24), FG (residues 116–132), and GH (residues 142–150) loops. The M20 loop is located directly over the active site, protecting it from the solvent, and is involved in the regulation of the active site [37]. The M20 loop is found in three conformations which are named as the open, occluded, and closed states [37, 40]. In our structural analysis, we have used the structure (PDB ID: 1rx2) that has the closed M20 loop conformation [37]. For each of the eleven mutants listed in Figure 2F as well as the wild type DHFR, we compiled 210 ns long MD simulations for both the DHFR/NADPH/DHF (green in Figure 2B) and the DHFR/NADPH/trimethoprim complexes (Materials and Methods) [41].

We have closely monitored the WT and all 11 single mutant sets of MD trajectories corresponding to those listed in Figure 2F to decipher the molecular mechanisms that lead to trimethoprim resistance. We note that while these mutations are observed with various frequencies in the morbidostat trajectories as displayed in Figure 1D, nine of them appeared as the first coding region mutation. Besides, the changes in the dynamics of the system due to resistance-conferring mutations are usually subtle. In particular, the effect on trimethoprim binding is indistinguishable in all DHFR/NADPH/trimethoprim complex simulations. This is expected since the free energy difference implied by the *K*_*i*_ changes reported in Figure 2F even in the most extreme case (∼30 fold increase for R98P) is predicted to be ∼2 kcal/mol. Such energy changes are often intractable in a conventional MD simulation with typical fluctuations occurring on the order of *RT* ≈ 0.6 kcal/mol. Nevertheless, it is important to note that despite the small differences in free energy, the local structural changes may be accommodated by entropy-enthalpy compensation as we have shown previously for the L28R mutant by isothermal titration calorimetry measurements [41]. This phenomena is explained by the utility of an interfacial water molecule as observed in the MD simulations [41]. Similarly, the MD simulations of the DHFR/NADPH/DHF complex do not implicate large dynamical changes in most of the MD trajectories. The three exceptions correspond to the most frequently observed first coding region mutations in the morbidostat, D27E, L28R, and W30R. Interestingly, in all three cases, distinct molecular strategies were utilized for rendering DHF more effective than trimethoprim (Figure 3).

**Figure 3.**
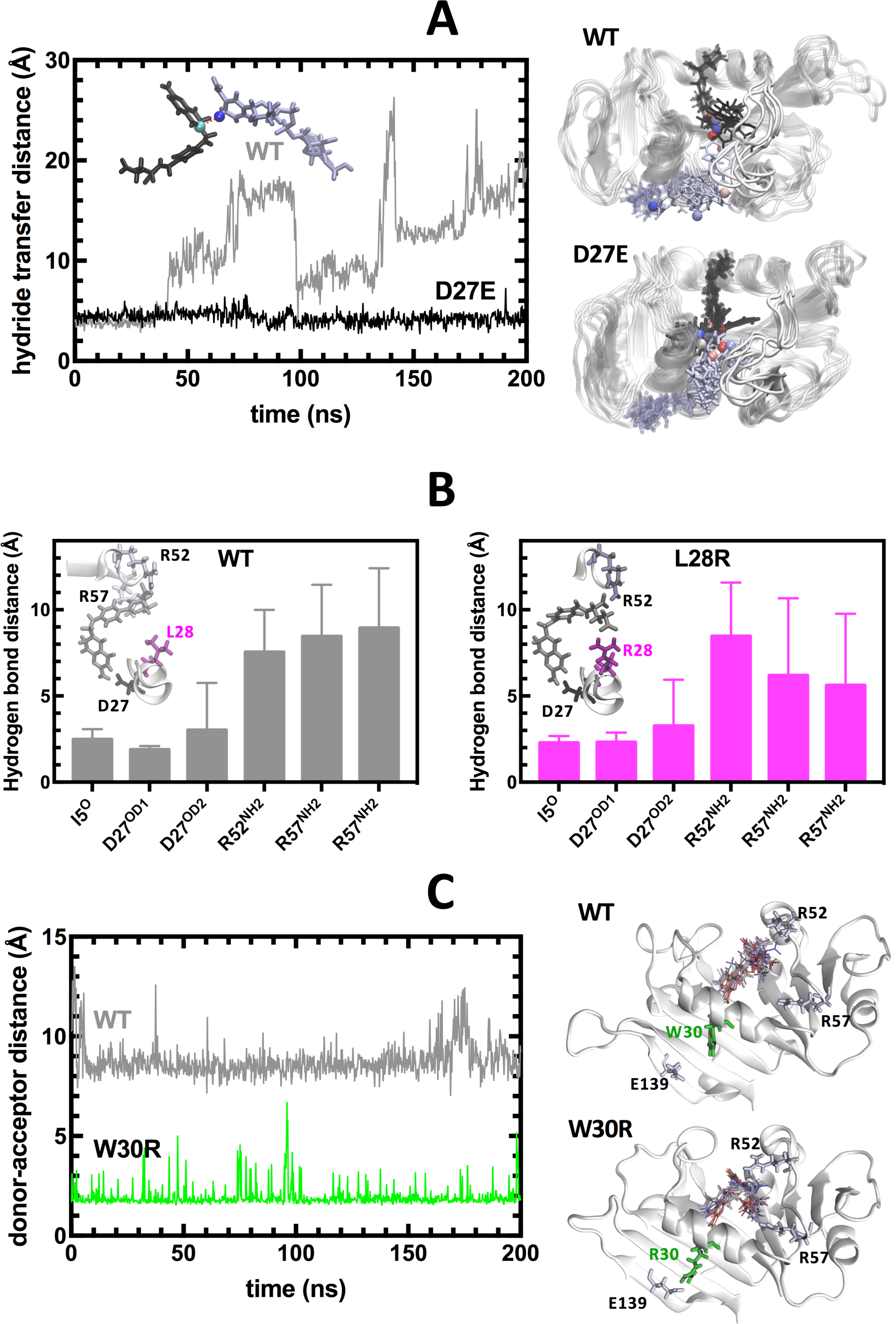
Molecular mechanisms operating in the DHF bound dynamics of DHFR for the three frequently observed DHFR mutations. **(A)** D27E replacement alters hydride transfer distance between the cofactor (NADPH) and the substrate (DHF). The measured distance is between the cyan and blue spheres shown in the inset for the crystal structure positioning of NADPH (black) and DHF, which is readily lost in the wild type structure as in all the other simulations of the single mutants except for D27E. Dynamical motions of NADPH and DHF are displayed on the right. **(B)** L28R mutations yields extra direct hydrogen bonds with DHF and stabilizes it in the binding pocket. The distance between the donors and acceptors of the hydrogen bonds originally present in the crystal structure is monitored throughout the MD trajectories with their averages and standard deviations displayed. While the original hydrogen bonds are lost in both the wild type and the L28R mutant, there are many new hydrogen bond donor sites on the R28 side chain, maintaining a dynamical hydrogen bonding ecology around the substrate. **(C)** W30R mutation releases the tension in the tight binding pocket by forming a salt bridge with E139. The distance between the E139 acceptor (O-group) and the closest heavy atom of residue 30 is plotted for the wild type and the W30R mutant. In the latter case a salt bridge is established between the side groups frequently, relaxing the tight binding pocket where the substrate resides. As shown on the right, DHF maintains a position between the stabilizing R52 and R57 side chains in the mutant while the contacts with R57 group is lost in the wild type.

In figure 3, we display resistance mechanisms for the D27E, L28R, and W30R mutations. Amongst the wild-type (WT) and all the single mutants we analyzed, the D27E mutant is the only one where the hydride transfer distance is kept at an optimal pre-catalytic range (Figure 3A). We note that in all mutations we studied, the M20 loop never leaves the closed conformation in favor of the occluded form which triggers the reduction of DHF into THF. Nevertheless, the longer side chain of the D27E mutant dynamically maintains the ligand at an optimal distance, keeping it ready for the hydride transfer once this rare event takes place, hence explaining the increase in *k*_*cat*_ for the D27E mutant (Figure 2F). On the other hand, the L28R mutation leads to the formation of extra hydrogen bonds between the enzyme and DHF, thus stabilizing its conformation [41]. In figure 3B, we display the average distance of hydrogen bonds formed between the enzyme and DHF. We find that while the pterin tail of DHF is permanently engaged in the binding pocket (as evidenced by the hydrogen bond distances to I5 and D27), the p-aminobenzoyl glutamate tail is mobile in the wild-type (WT) DHFR. In contrast, this mobility is significantly reduced in the L28R mutant due to the extra interactions provided by the side chain. Unlike D27E and L28R, the effect of W30R on the dynamics of DHF is indirect. In this case, the R30 side chain of the mutant forms a salt bridge with the side chain of E139 residing on the β sheet supporting the catalytic region (Figure 3C). The distance between the two residues is reduced from a baseline value of ∼8 Å to ∼2 Å. This interaction slightly opens the tight binding pocket so that the DHF p-aminobenzoyl glutamate tail motions are accommodated in the region between R52 and R57 residues, whereas the glutamate tail is more disordered and closer to R52 residue in the wild-type DHFR. Reduced interactions between the p-aminobenzoyl glutamate tail and the enzyme leads to weaker substrate binding and higher catalytic rate.

To summarize our experimental and computational findings thus far, we conclude that the effect of single DHFR mutations on trimethoprim binding is not definitive for survival. It is rather the small changes on the binding kinetics of the substrate (DHF) that provide the enzyme a small advantage that is utilized for bacterial survival. Furthermore, the changes in the DHF binding dynamics induced by single mutations are diverse. In the rest of the manuscript, we discuss the changes in the fitness landscapes due to the accumulation of multiple mutations using a library of combined mutants selected from a subset of those observed in the morbidostat trajectories.

### Trimethoprim-free enzymatic velocity of DHFR mutants correlates well with trimethoprim-free growth rates of *E. coli* mutants carrying corresponding DHFR mutations

Resistance-conferring mutations are rarely found in natural bacterial isolates and this observation is generally attributed to the fitness costs of resistance-conferring mutations. In the case of enzymes such as DHFR, where multiple resistance conferring mutations are sequentially fixed, it is not clear how that many mutations can be tolerated and yet sufficient enzymatic activity is maintained for organismal survival. To address this question, we selected six of the mutations listed in Figure 2F (P21L, A26T, L28R, W30G, W30R, and I94L) and synthesized a DHFR mutant library where we had all 48 (3^1^x2^4^) possible combinations of these mutations. We purified and characterized all of the mutant DHFR enzymes as previously described (Table S3). Next, we measured growth rates of the *E. coli* mutant library (Figure 4A) that carry the same DHFR mutations in various conditions (different temperature, different glucose concentrations, and different casamino acids concentrations) (Figure 4B-F). We found that enzymatic activity of DHFR mutants in the absence of trimethoprim (*V*_*0*_, equation 1), calculated at saturating [DHF], correlated well with the trimethoprim-free growth rates of *E. coli* mutants with corresponding DHFR mutations (*r* = 0.46-0.58, *p* < 10^−3^, Pearson Correlation Test). The correlations between growth rates and other biochemical parameters such as *k*_*cat*_ or *k*_*cat*_/*K*_*m*_ were less significant (for *k*_*cat*_: *(r* = 0.33, *p* < 10^−3^); for *k*_*cat*_/*K*_*m*_: *(r* = 0.06, *p* < 10^−3^), Pearson Correlation). We note that the 12.5μM DHF concentration is in good agreement with the previously measured *in vivo* DHF concentrations in which both reduced and oxidized species of folate concentrations were in the range of ∼10 μM [42]. These experiments and the resulting analysis suggested that *V*_*0*_, the substrate reduction rate of DHFR in the absence of trimethoprim, is a good predictor of bacterial fitness, particularly when limited nutrients are provided to bacterial populations (i.e., minimal media supplemented with 0.4% glucose) and bacterial cells are grown in the absence of trimethoprim.

**Figure 4:**
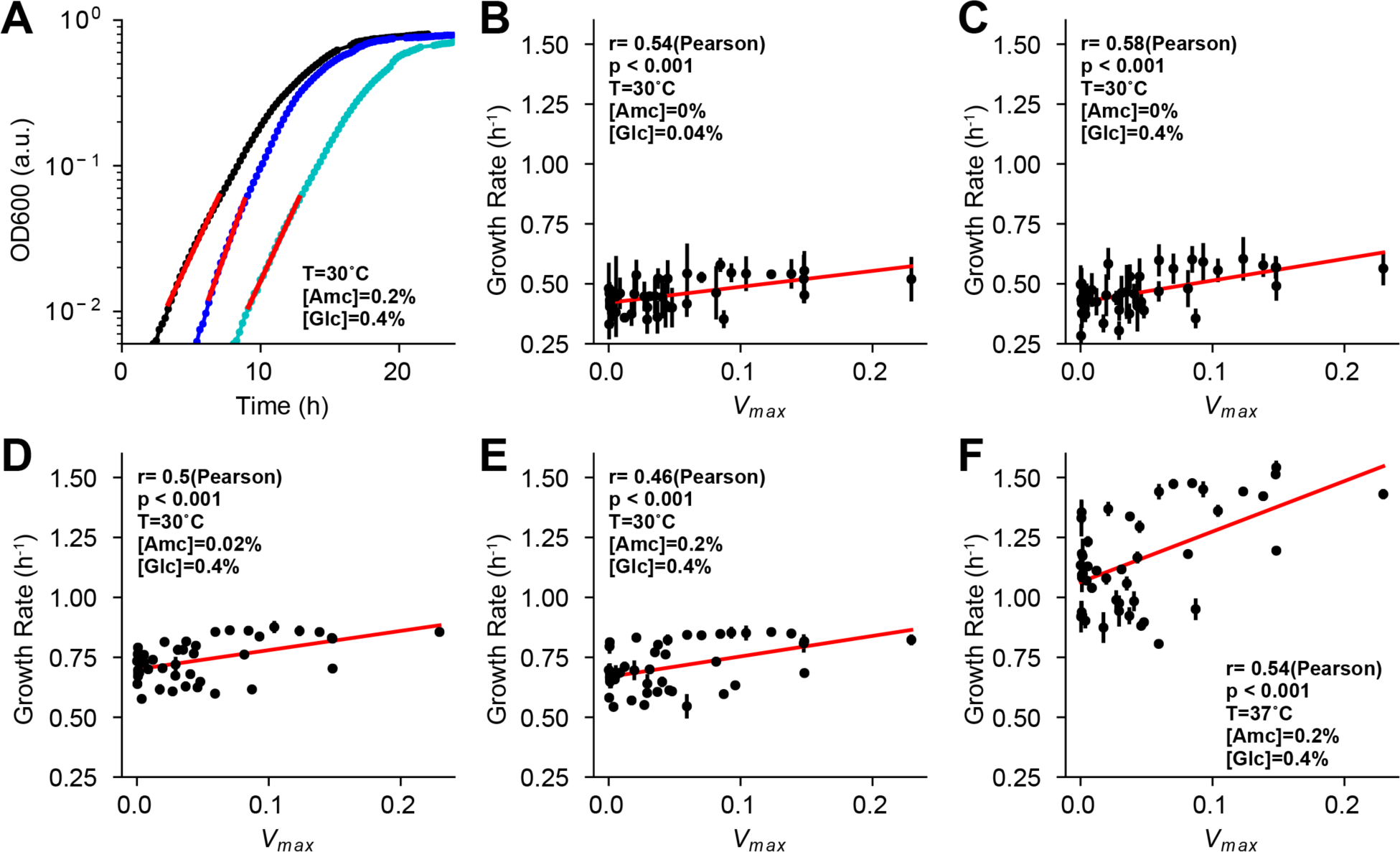
Bacterial growth rates correlate with DHFR’s enzymatic activity. **A)** Growth rates (*μ*) of E. coli cells with DHFR mutations are calculated by fitting an exponential growth function; OD(t) = OD(0). e^μ.t^, to the cell density (OD600) readings. **B-F)** Mean growth rate values (± standard deviation) of all mutations are measured for different M9 minimal media compositions and temperature (*T*). Correlation between *V*_*0*_ and growth rate is calculated using Pearson Correlation test. *r*: correlation coefficient, *p*: significance. [Amc] stands for amicase concentration; [Glc] stands for glucose concentration.

### Combined effects of resistance-conferring mutations deviate from fitness values predicted by Bliss Additivity

In order to qualitatively understand the evolutionary trade-offs in DHFR evolution, we plotted *V*_*0*_ values against the corresponding *K*_*i*_ values for DHFR mutants. Interestingly, *V*_*0*_ values exhibited a bifurcation in this geometric representation (Figure 5A). DHFR mutants either had enzymatic activities comparable to their wild type ancestor or significantly lost their enzymatic activities, displaying almost no activity. Interestingly, all of the mutants that were funneled into the highly decreased enzymatic activity regimen carried the P21L mutation (Figure 5A, red triangles and circles). In addition, none of the mutants that were detected in the morbidostat (Figure 5A, grey and red circles) had *V*_*0*_ values lower than four percent of the wild type *V*_*0*_ (Figure 5A, horizontal dashed line). We note that all of the DHFR alleles observed in the morbidostat appeared in the background of a promoter mutation that increases DHFR amount by 10-20 fold (Figure 1B, insert). Therefore, all the observed mutants in the morbidostat are predicted to have DHFR activity equivalent to 40-80 percent of the wild type DHFR (*V*_*0*_).

**Figure 5:**
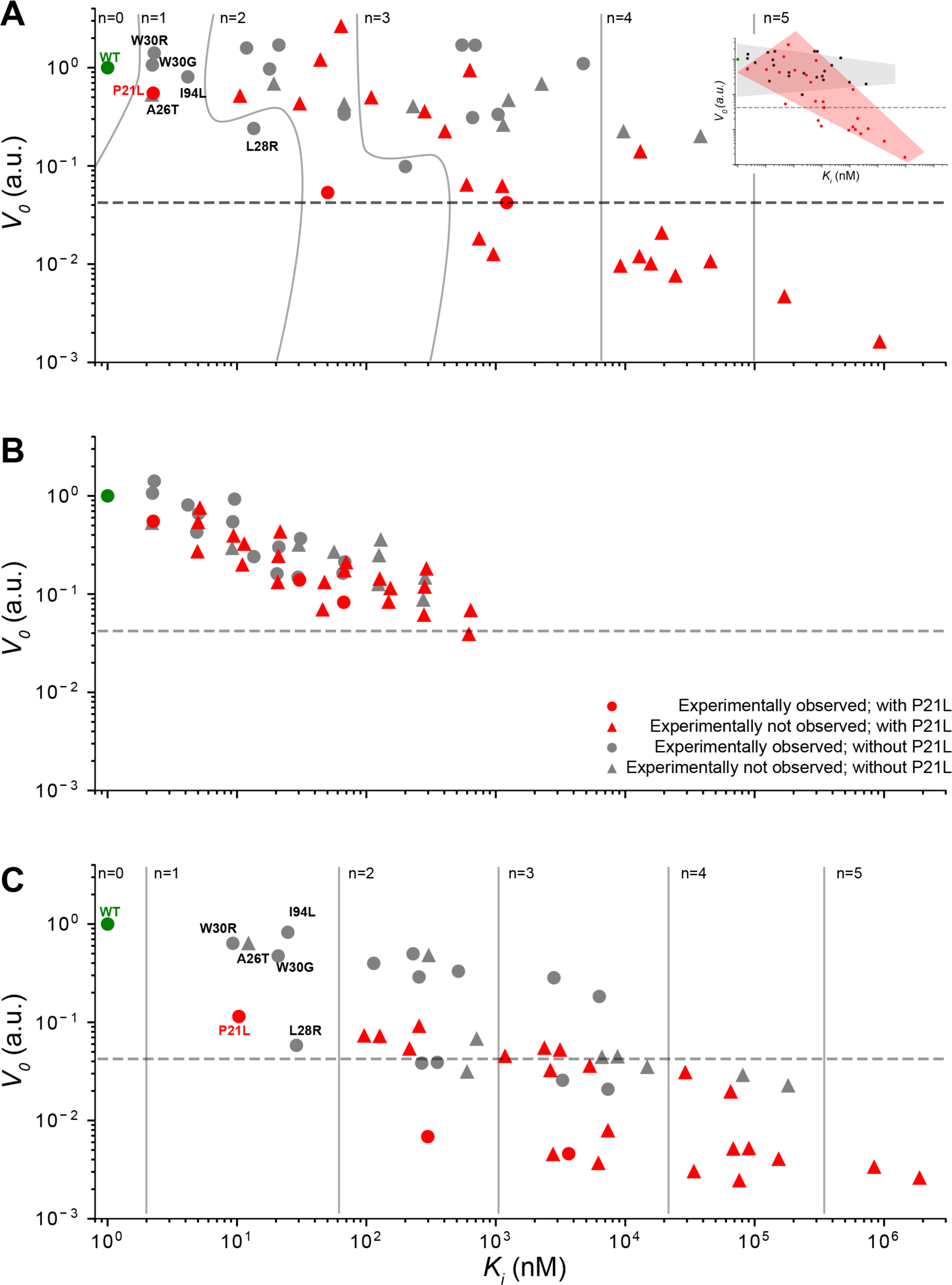
Combined effects of resistance-conferring mutations deviate from fitness values predicted by Bliss Additivity model. **A)** *V*_*0*_ vs *K*_*i*_ values of the 48 DHFR mutants are plotted. Curved and straight lines are used to separate mutants with different number of mutations. Horizontal dashed line shows the minimum *V*_*0*_ value for a DHFR mutant that was observed in the morbidostat experiment. Red markers show mutants with P21L mutation. Gray markers show mutants without P21L mutation. Circle markers show mutants that are observed in evolution experiments. (Insert) *V*_*0*_ values bifurcate depending on the presence of P21L mutation. **B)** Predicted *V*_*0*_ and *K*_*i*_ values for multiple DHFR mutants by Bliss Independence model using the *V*_*0*_ and *K*_*i*_ values measured for DHFR variant with single mutations (relative to the wild-type DHFR). These predictions significantly deviate from experimental observations (both for *V*_*0*_, and for *K*_*i*_ (Student t-test, p<10^−3^). This model under-predicts *K*_*i*_ values by a factor of 0.27 ±0.35 and over-predicts *V*_*0*_ values by 3.34 ± 0.35 (Mean ± standard deviation; Figure S1, Table S3). **C)** Predicted *V*_*0*_ and *K*_*i*_ values for multiple DHFR mutants by Bliss Independence model using the (geometric) mean effects of single mutations on all possible genetic backgrounds (Table S4). This model over-predicts *K*_*i*_ values by a factor of 6 ±3.96 and under-predicts *V*_*0*_ values by 0.35 ± 0.39 (Mean ± standard deviation; Figure S1, Table S3). The bifurcation observed in panel A disappears in both analysis summarized in panels B and C.

In order to test the existence of epistatic interactions among DHFR mutations, we asked whether the *K*_*i*_ and *V*_*0*_ values deviated from the *K*_*i*_ and *V*_*0*_ values predicted by using an additive model, assuming Bliss independence between the effects of the mutations [43]. According to Bliss independence, effects of multiple mutations should simply add up to the sum of the individual effects of mutations. However, as shown in Figure 5B, when the individual effects of six single mutations on the wild type DHFR are used to calculate *K*_*i*_ and *V*_*0*_ values using the Bliss additivity [43], the predicted *K*_*i*_ and *V*_*0*_ values were significantly different from the experimentally measured ones (Student t-test, p<10^−3^; Figure S1, Table S3). We also found that the predicted *V*_*0*_ values did not display a bifurcation and steadily declined as the number of DHFR mutations increased. We also found that the predicted *K*_*i*_ values were not as large as the experimentally measured values (Figure 5 A-B). When we instead utilized the mean effects of single mutations on all possible genetic backgrounds in our mutant library (Figure 6), we were able to better estimate the *K*_*i*_ values (Figure 5C). However, the bifurcation we observed in Figure 5A disappeared and several of the mutants had lower predicted *V*_*0*_ values compared to the experimentally measured ones. These observations clearly suggested the existence of epistasis (deviation from additivity) among the six DHFR mutations we studied. The effects of DHFR mutations seemed to be context dependent and recovering the fitness of DHFR mutants with multiple mutations would at least require mean fitness effects of mutations calculated on several different genetic backgrounds.

**Figure 6:**
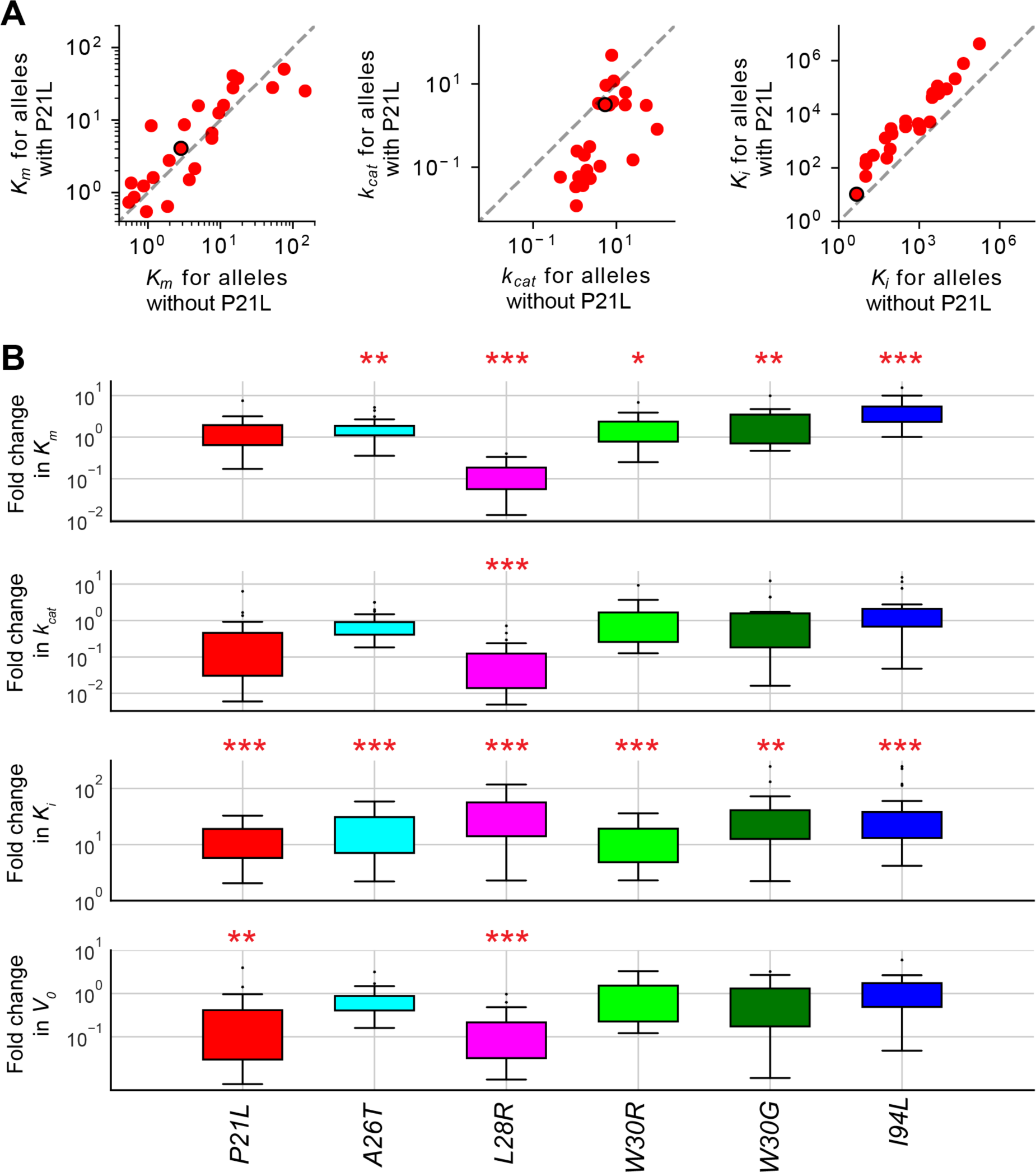
Mean effects of DHFR mutations in catalytic activity and trimethoprim binding. **A)** Each marker in upper panels show fitness changes when a mutant acquires P21L mutation. x axis shows *K*_*m*_, *k*_*cat*_ and *K*_*i*_ values of mutant alleles without P21L mutation and y axis shows the values mutant alleles with P21L mutation. For instance, the black encircled points has the *K*_*m*_, *k*_*cat*_ or *K*_*i*_ value of WT on x axis and corresponding values for P21L on y axis. **B)** Fold change effects when each single mutant is added on top of all other genotypes. Student’s t-test (two tailed) is used to quantify significance of *K*_*m*_, *k*_*cat*_ and *K*_*i*_, *V*_*0*_ changes relative to the wild type DHFR (*: p<0.05; **: p<0.01; ***: p<0.001).

### Effects of mutations on the catalytic power of DHFR were largely context dependent due to epistasis between mutations

We calculated (geometric) mean phenotypic effects of individual DHFR mutations on *K*_*m*_, *k*_*cat*_, and *K*_*i*_ and *V*_*0*_ values (Table S3). Briefly, for every single amino acid replacement in DHFR, we divided the DHFR mutant library into two groups depending on whether they have a particular mutation (i.e. P21L, Figure 6A) and compared the *K*_*m*_, *k*_*cat*_, *K*_*i*_, and *V*_*0*_ values of the two groups. Thus, we were able to calculate mean fold changes in *K*_*m*_, *k*_*cat*_, *K*_*i*_, and *V*_*0*_ values due to a single mutation as shown in Figure 6B. This analysis clearly showed that all six of the mutations we analyzed increased the *K*_*i*_ values on all possible genetic backgrounds explaining their resistance-conferring effects. Similarly, all of the mutations except L28R and P21L, significantly decreased substrate affinity (increased *K*_*m*_). As discussed before, the L28R mutation increased substrate affinity of DHFR. The mean effect of P21L mutation on *K*_*m*_ was not statistically significant. However, although L28R decreased *k*_*cat*_ values on almost all possible genetic backgrounds, the rest of the mutations did not have statistically significant effects on *k*_*cat*_ values. The large variations in the mean effects of these mutations on *k*_*cat*_ values suggested that the effects of mutations on the catalytic power of DHFR were largely context dependent due to epistasis between mutations.

### Epistasis between resistance-conferring DHFR mutations is high-order for substrate binding and catalysis (*k*_*cat*_ and *K*_*m*_) but first-order for drug binding (*K*_*i*_)

We quantified epistatic interactions between the six DHFR mutations (P21L, A26T, L28R, W30G, W30R, and I94L) we studied by utilizing a linear regression model (Materials and Methods) [44]). Briefly, we attempted to recover fitness values of all DHFR alleles using epistatic terms between mutations. In a biological system, if the epistasis between mutations is large, it is difficult to recover fitness values for genotypes with *n* mutations by using up to *m*^th^ order epistatic terms (*m* < *n*). However, if epistatic interactions are less prevalent, predicting fitness of genotypes by using up to *m*^th^ order epistatic terms (m < n) becomes more feasible. As shown in Figure 7A, we were able to adequately predict *K*_*i*_ values for all DHFR mutants with up to five mutations by using only the first order epistasis terms (∼10-20% residual error). The extra information we gain from using higher order epistatic terms was relatively small (Figure 7B) indicating that measuring the *K*_*i*_ values of single DHFR mutations and first order epistatic terms (mathematically equivalent to mean effect of mutations) will mostly be sufficient to predict *K*_*i*_ values of DHFR mutants with any combination of the six DHFR mutations we studied. This analysis is consistent with our findings summarized in Figures 5 and 6. On the contrary, predicting *k*_*cat*_ and *K*_*m*_ values of DHFR mutants (with multiple mutations) by using epistatic terms was relatively more challenging due to high-order epistasis. For both *k*_*cat*_ and *K*_*m*_, in order to obtain a prediction power comparable with what we had for *K*_*i*_, we needed to use at least up to third order epistatic terms and yet there was a big variance in the prediction performance (Figure 7B). This suggested that the effects of the mutations on DHFR’s catalytic activity were highly context dependent which make fitness landscape of DHFR rugged [15]. We conclude that the epistasis between resistance-conferring mutations is high-order for *k*_*cat*_ and *K*_*m*_ but first-order for *K*_*i*_ values. Since DHFR fitness in trimethoprim containing environment is a convoluted function of all *k*_*cat*_, *K*_*m*_, and *K*_*i*_ values, evolution of trimethoprim resistance in the adaptive landscape is mostly unpredictable mainly because of high-order epistatic interactions in catalytic power of DHFR (*k*_*cat*_ and *K*_*m*_).

**Figure 7:**
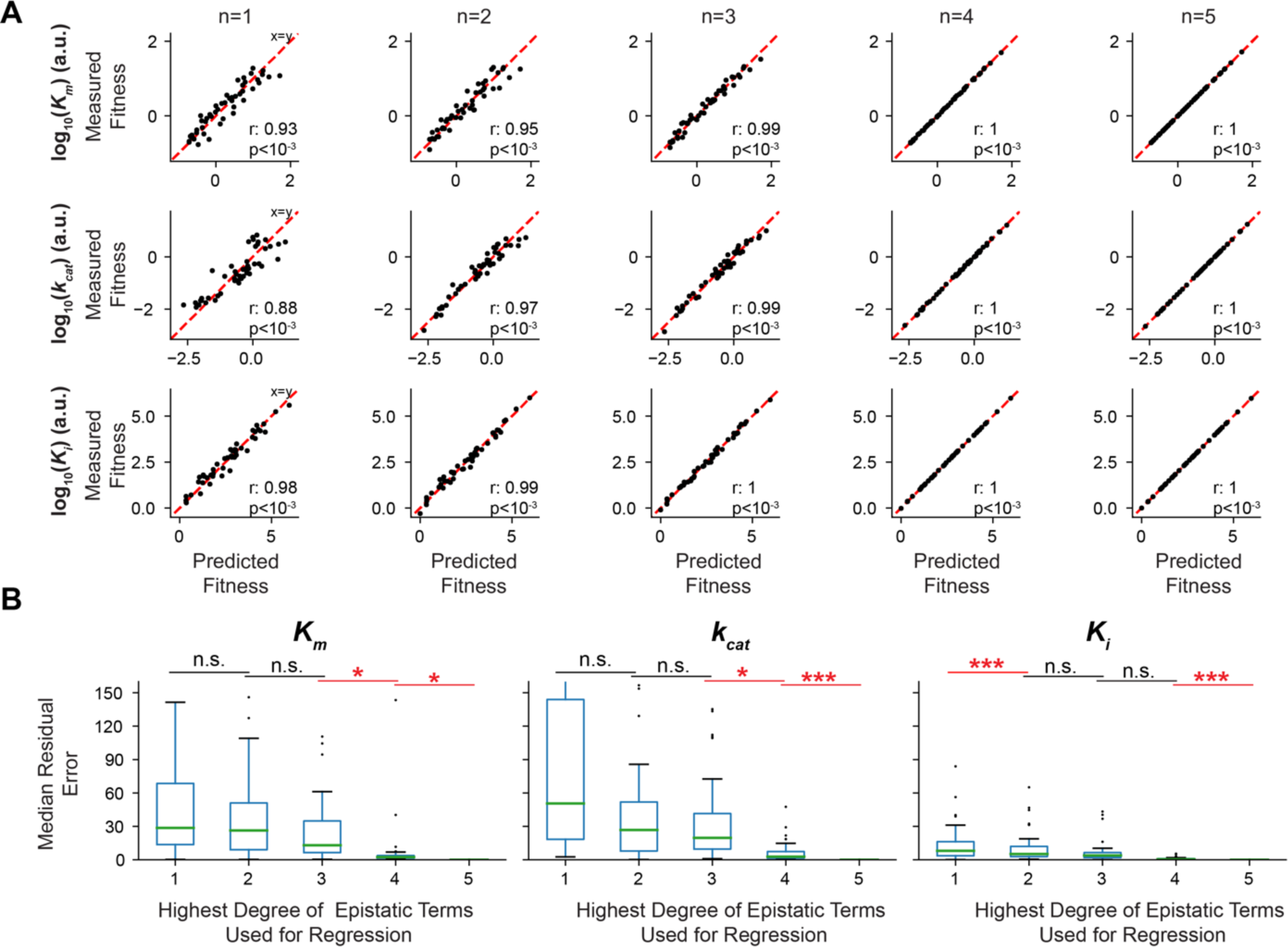
Epistasis between resistance-conferring DHFR mutations is high-order for substrate binding and catalysis (*k*_*cat*_ and *K*_*m*_). **A)** A linear regression model is used to predict fitness information stored in epistatic terms with increasing orders. Correlations between predicted fitness values of all genotypes using *n*^*th*^ order epistatic terms and the measured fitness values are calculated. **B)** Median residual errors for predicted fitness values as function of degree of epistatic terms used in regression. First order epistatic terms are sufficient to recover experimental *K*_*i*_ values with ∼10-20% residual error. However, at least second and third order epistatic terms are required to recover experimental *K*_*m*_ and *k*_*cat*_ values with with ∼10-20% residual error.

### MD simulations demonstrate the context dependent effects of DHFR mutations at the molecular level

Epistatic interactions in biological systems are common and were previously reported by several researchers. However, in most cases, the molecular basis of epistasis was not sufficiently explained [6, 26]. To study molecular basis of epistasis between resistance-conferring DHFR mutations, we utilized MD simulations [41]. Since our biochemical analysis and epistasis calculations suggested that the epistasis was largely due to substrate binding and catalysis, we performed MD simulations for the substrate-bound conformation of DHFR (Methods). We carried out MD simulations for a subset of DHFR alleles including all combinations of the mutations A26T, L28R and I94L. In addition, we traced the effect of adding P21L mutation to some of these mutants in order to understand how adding the P21L mutation drastically reduces enzymatic efficiency (Figures 5 and 6). Amongst these, L28R is frequently observed as the first coding region mutation in the morbidostat while A26T and I94L are observed later in evolution experiments (Table S5).

We demonstrate the context dependence of the observed dynamics by focusing on four specific examples involving double mutations in Figure 8. We traced the signature hydrogen bonds between the enzyme and the substrate (Figure 8) and found that hydrogen bonds between the I5 and D27 side chains in the studied mutants were always close to their native values in the wild type DHFR. However, the hydrogen bonds between the R52 and R57 side chains and DHF showed significant variations (displayed in figure 8, averaged over the last 100 ns of the trajectories.) For the single mutants, we do not find any significant dynamical changes in the MD trajectories for P21L and A26T mutations. We note that the common reduction in the *k*_*cat*_ value due to the P21L mutation (Figure 2F) implied that the effect of this mutation is mainly in the dynamics of the catalytic M20 loop, whose dynamics is on the time scale of seconds and is therefore not within the sub-microsecond observation window of our MD simulations. Meanwhile, the I94L mutant completely loses interactions with the R57 side chain since the slight change in the isomerization of the side chain leads to more prolonged interactions with the aromatic ring of DHF, distorting the tight binding pocket. As a result, the R57 side chain flips out of the pocket to the other side of the helix spanning residues 25-37 (figure 8A).

**Figure 8.**
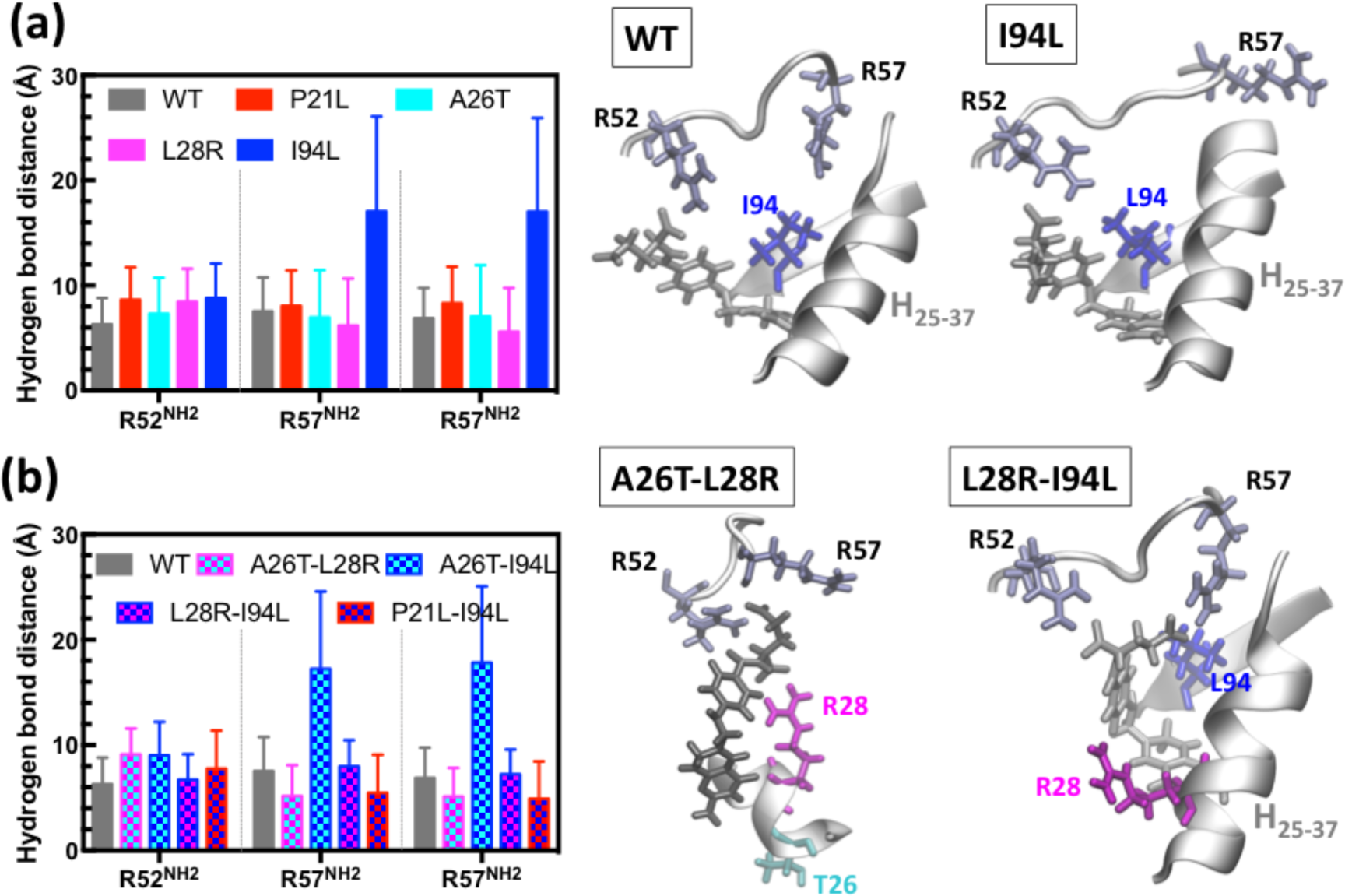
Epistasis between resistance-conferring DHFR mutations are largely due to interactions of the mutated enzyme with the p-aminobenzoyl glutamate tail of DHF. (A) The I94L mutation exacerbates substrate binding of DHFR by altering tight interactions with the p-aminobenzoyl glutamate tail of DHF in the binding pocket, allowing the R57 side chain to flip out. (B) L28R mutation is a highly epistatic mutation; together with either A26T or I94L, the L28R further stabilizes the substrate in the pocket. Note that the P21L-I94L double mutation also rescues the negative effect of I94L as traced through the hydrogen bond distances.

As was previously described in figure 3B, L28R mutation leads to the formation of extra hydrogen bonds with DHF. We found that together with A26T, this effect becomes even stronger, fixing the position of DHF to the space between R52 and R57 residues (figure 8B). Thus, while the A26T mutation alone causes subtle structural changes in our MD simulations, together with L28R, it benefits from a synergistic effect on DHF binding, with the polar side chain further stabilizing the network of hydrogen bonds in the pocket. The L28R mutation has a similar synergistic effect on the I94L mutation. Despite the tendency of the I94L mutant to interact strongly with the aromatic part of DHF, the binding pocket is not as easy to distort due to the presence of R28 interactions with the substrate, leading to a stabilized ligand (DHF). We note that addition of A26T to the I94L mutation does not have the same synergistic effect as expected by the outlined mechanism of action. Interestingly, although P21L mutation mostly impairs catalytic activity of DHFR, the P21L mutation rescues I94L mutant. In this case, the more flexible L21 allows distortions of the tight binding pocket without letting the R57 side chain to flip out (not displayed). We note that these mutations significantly decrease the binding propensity of the inhibitor, as measured by the *K*_*i*_ values listed in Table S2. DHF escapes this fate due to the extra interactions of the larger ligand with the side chains of the enzyme. Running longer MD simulations for all possible combinations of DHFR mutations was beyond our computational capacity but even the analysis of this small subset of DHFR mutants demonstrated the context dependent effects of DHFR mutations at the molecular level.

On the other hand, we do not observe significant structural fluctuations in DHFR upon trimethoprim binding unless more than two mutations are accumulated. With the introduction of a third mutation, the dynamics of the DHFR is significantly altered, with amplified motions observed in the loops. In Figure S2A, we display the root mean square fluctuations (RMSF) as mutations are accumulated. With the triplet A26T-L28R-I94L large motions in new regions are observed along with a substantial increase in the amount of fluctuations; the effect is magnified as more mutations are accrued. In fact, the RMSF of multiple mutants are well correlated with log *K*_*i*_ values as displayed in Figure S2B, (multiple mutants: *r* = 0.70, p < 0.01; all cases including WT: *r* = 0.60, *p* < 0.01; Pearson Correlation). Thus, the effect of decreased inhibitor binding affinity is significant for reinforcing resistance in higher number of mutants, while the first two mutations are more effective in the catalytic activity.

### Promoter mutations compensate detrimental effects of several mutations and largely increases number of plausible evolutionary trajectories

Evolution of trimethoprim resistance is a random search for mutational trajectories that lead to the resistant DHFR genotypes without sacrificing catalytic activity. We ran computer simulations to visualize and quantify plausible evolutionary trajectories leading to trimethoprim resistance. As demonstrated in Figure 9, for every DHFR allele, we calculated DHFR activity (*V*) as a function of trimethoprim concentration. In Figure 9, DHFR mutants are represented as cylindrical pillars with heights proportional to trimethoprim concentrations necessary to reduce mutated DHFR enzymes’ activities down to 50% of *V*_*0*_ (*V*_*0*_^*WT*^) for the wild type DHFR. Colored filled circles on the upper surface of the cylinder represent DHFR mutations. We note that this landscape dynamically changes as we increase trimethoprim concentrations used in our calculations. In these calculations (Equation 1), we used a saturating dihydrofolate (DHF) concentration (12.5 μM) which is in the physiological range and we assumed a ten-fold increase in DHFR expression due to the promoter mutation (Figure 1B). Alleles are grouped according to the number of mutations they have. We then ran stochastic simulations where we consider the DHFR sequence as a lattice and allow DHFR to acquire mutations as trimethoprim dosage is gradually increased. All simulations start from the wild type DHFR allele and the activities of all DHFR alleles are calculated at every trimethoprim concentration. In these simulations, we assume that any DHFR mutant that has activity (*V*) less than half of the wild type DHFR activity (*V*_*0*_^*WT*^, no trimethoprim) goes extinct unless they acquire a beneficial mutation. In our simulations, we allow DHFR to obtain or lose one of the seven mutations (promoter, P21L, A26T, L28R, W30G, W30R, and I94L) if activity of the mutant is about to drop below half of *V*_*0*_^*WT*^. Any of these mutations can be added, converted (W30R → W30G, W30G →W30R) or reverted (e.g. L21 mutant to P21). As shown in Figure 9, we observed several genetic trajectories that arrive at local or global maxima. We repeated these simulations 10^6^ times and quantified relative abundance of mutational trajectories (Figure 9 and Table S6).

**Figure 9:**
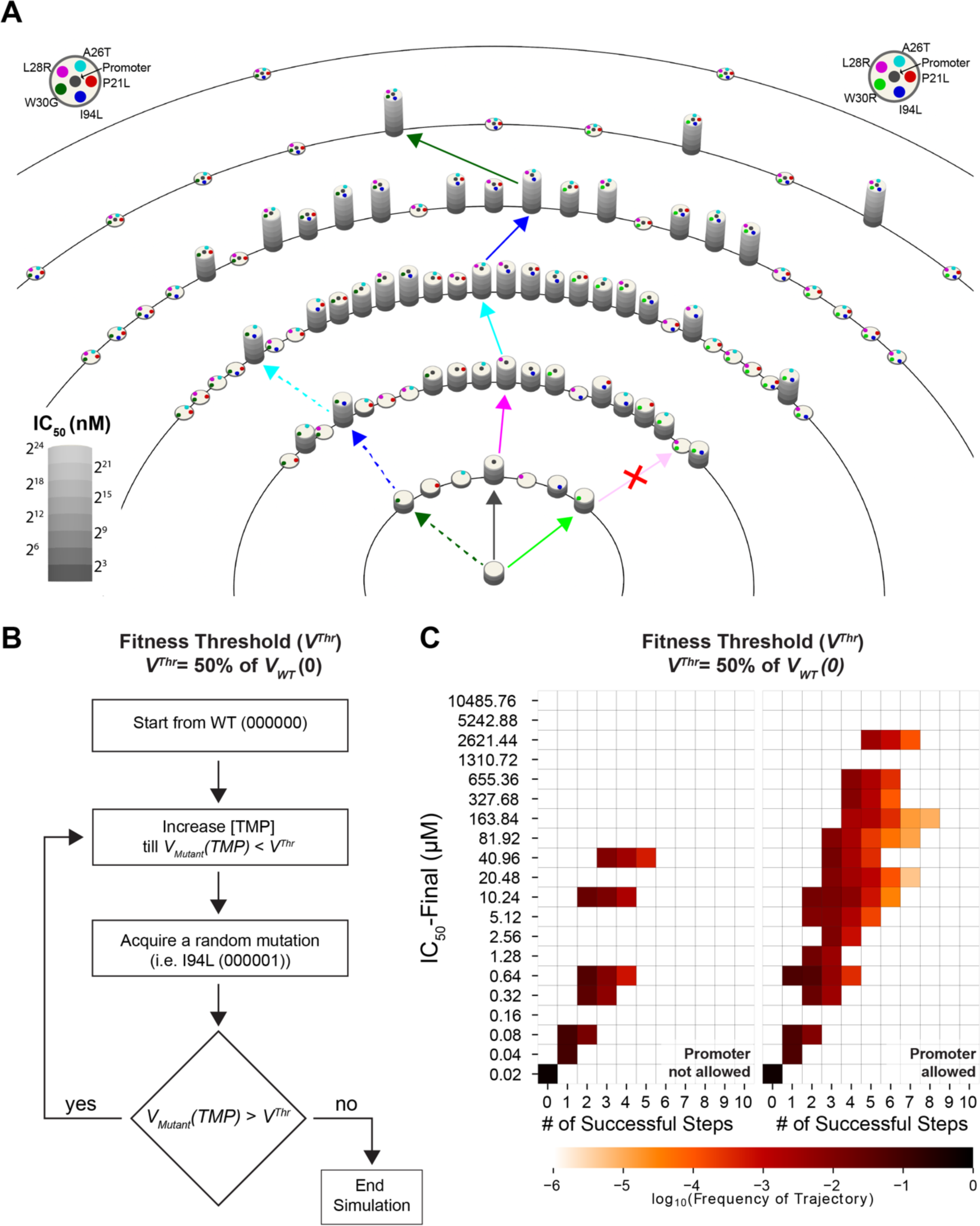
Simulated evolutionary trajectories leading to trimethoprim resistance. **A)** DHFR alleles are represented as cylindrical pillars. Atop of pillars, colored filled circles are used to show DHFR mutation. Heights of the cylinders correspond to trimethoprim concentrations required to reduce the activity of mutant DHFR enzymes down to half of the *V*_*0*_ for the wild type DHFR (*V*_*0*_^*WT*^). Note that several pillars have zero height because their activities never exceed half of *V*_*0*_^*WT*^ even in the absence of trimethoprim. The trajectory represented with solid arrowed lines is one of the shortest and most common pathways leading to global maximum of the adaptive landscape. The trajectory represented with dashed arrowed lines lead to a local maximum of the adaptive landscape if the promoter mutation is not allowed. **B)** Schematics summarizing the algorithm used in simulations. **C)** Simulations analysis summarized in heat maps. In simulations where the promoter mutation is not allowed (left), trajectories are shorter compared to the trajectories where the promoter mutation is allowed (right). If the promoter mutation is allowed, an increased number of trajectories lead to adaptive peaks with higher trimethoprim resistance levels.

Mutational trajectories that lead to high trimethoprim resistance peaks typically accumulated up to five mutations and the majority of these trajectories reached to the fitness peaks in five to seven genetic steps. Several viable trajectories included more than five mutational steps mainly because reverting the P21L mutation back to wild-type (L21P) significantly improved DHFR fitness in several genetic backgrounds. We then ranked all of the genetic trajectories that reach to high trimethoprim resistance by taking the least possible number of steps and calculated the likelihood of each mutation in the adaptive landscape (Table S6). We have also repeated these simulations using lower fitness thresholds (i.e. 1% of V_0_ for the wild type DHFR) and showed that number and length of evolutionary trajectories that reach to fitness peaks drastically increase if minimum fitness thresholds are assumed to be lower (Figure S3).

Finally, we computationally tested the effect of promoter mutations in DHFR evolution (Figure 9C). To do this, we ran simulations where all of the DHFR alleles with promoter mutations were eliminated and we compared these simulations with those that allow the promoter mutation. We found that number of plausible mutational trajectories that lead to trimethoprim resistant genotypes significantly diminishes if the promoter mutation is not allowed (Figure 9C). When promoter is not allowed, only 1.289 ± 0.005% of the simulated trajectories reach to genotypes that survived in 32 μM trimethoprim which is considered as resistant in clinical microbiology laboratories. There are only 60 unique trajectories which acquired one or more DHFR mutations and increased trimethoprim resistance. However, when promoter mutation is allowed, 5.592 ± 0.026% of the simulated trajectories reach to genotypes that survived in trimethoprim concentrations between 32 μM and ∼2.58mM. In this case, 2573 unique trajectories acquired one or more DHFR mutations and increased trimethoprim resistance. This effect is mainly due to elimination of half of the possible genetic combinations between the six resistance-conferring mutations we studied and also elimination of the compensatory effect of the promoter mutation. Thus, number and length of plausible evolutionary trajectories, as well as the maximum possible trimethoprim resistance significantly diminish in the absence of the promoter mutation. Therefore, in the absence of promoter mutation, DHFR evolution becomes more predictable. As a result, being able to target the promoter mutation with one of the novel gene editing methods together with a mutant-specific drug that specifically inhibits a mutation such as L28R, that is a synergistic mutation, might significantly slow down evolution of trimethoprim resistance. We note that eliminating the promoter mutation or the L28R mutation does not exclude other evolutionary solutions such as acquiring other resistance conferring mutations listed in Figure 2F, gene duplication, and acquiring other promoter mutations.

We conclude that, although expected to be random, the first plausible mutation in DHFR evolution is expected to be one of the promoter, W30R, or W30G mutations. Indeed, the c-35t and W30R mutations were previously found in clinically isolated *E. coli* strains [45]. Due to epistatic interactions, evolutionary trajectories become more constrained after acquiring the second and third mutations. However, the promoter mutation makes the adaptive landscape of DHFR less predictable by compensating for diminished catalytic activities of resistance-conferring DHFR mutation(s).

## Discussion

DHFR is a ubiquitous enzyme commonly used as a drug target in antibacterial, anticancer, and antimalarial therapies [21]. Developing a better understanding of the evolution of drug resistance through sequential accumulation of DHFR mutations is therefore an important scientific task to help improve drug therapies. Our experimental findings and computational analyses demonstrate that DHFR is a highly evolvable enzyme that can maintain its catalytic activity while accumulating multiple resistance-conferring mutations. Throughout the evolution of trimethoprim resistance in *E. coli*, DHFR can accumulate mutations in at least ten residues and four different promoter positions. In addition, amplification of chromosomal regions spanning the *folA* gene that encodes for DHFR is rarely observed [26]. Experimental and computational analysis of six of these mutations demonstrate the prevalence of epistatic interactions between them which imply the ruggedness of the adaptive landscape that lead to trimethoprim resistance. Epistasis between resistance-conferring mutations in *E. coli* DHFR and PfDHFR was previously reported and quantified by engineering all possible combinations of a small number of resistance-conferring mutations [15, 22]. A similar analysis was also done for a beta-lactamase gene in the landmark study of Weinreich and Hartl [6]. These studies mainly utilized bacterial growth assays to quantify fitness effects of mutations and assessed the predictability for evolution of resistance. In another landmark study by Lunzer *et al.,* where they systematically studied effects of amino acid changes in isopropylmalate dehydrogenase’s coenzyme choice, they demonstrated that each amino acid additively contributed to the function of isopropylmalate dehydrogenase’s enzymatic function, and that the epistasis comes from non-linearities in the fitness [46]. Conversely, in this study, by utilizing both biochemical assays and growth rate measurements, we deconvolved epistasis between resistance-conferring mutations and demonstrated that epistasis was largely due to changes in catalytic activity of the mutant DHFR enzymes rather than nonlinearity in bacterial fitness. We also showed that epistatic interactions and the compensatory effects of promoter mutations significantly diminish our ability to predict DHFR evolution in the presence of trimethoprim induced selection.

In a recent study, Rodrigues *et al.* investigated epistasis between three of the mutations we studied (P21L, L28R, and W30R) and developed an elegant framework to predict fitness of *E. coli* strains carrying DHFR alleles with combinations of these three mutations by using biophysical properties of DHFR mutations [7]. However, because of the low number of possible combinations (2^3^) of DHFR mutations they studied, they were not able to observe the P21L-caused bifurcation in the fitness landscape we report here (Figure 5). Therefore, for a larger set of combinations of DHFR mutations that include the P21L, fitness prediction of DHFR alleles will naturally be more difficult. Using the available biochemical fitness values we have, we were able to identify partial correlation between catalytic power and bacterial growth rates of DHFR mutants. However, we were not able to demonstrate a direct correlation between trimethoprim resistance and biochemical parameters we measured. We note that predicting trimethoprim resistance levels might be possible by using extra biophysical parameters such as thermal stability and abundance of DHFR mutants as was demonstrated by Rodrigues *et al.* [7].

Our analysis suggests that although predicting DHFR evolution is a difficult task, it might still be possible to steer evolution of trimethoprim resistance towards clinically less challenging phenotypes. Among all the mutations we studied, promoter and L28R mutations can potentially be targeted to reduce the number of plausible evolutionary trajectories and trimethoprim resistance. For example, being able to specifically target the promoter mutation by utilizing one of the novel gene editing tools will substantially decrease both the number of accessible trajectories and maximum resistance levels (Figure 9) [47]. Also, since the L28R mutation has a distinct molecular mechanism that increases its relative preference for the substrate over the drug molecules (Figure 3), it might be possible to design L28R-specific DHFR inhibitors that will mimic DHF without losing its specificity against bacterial DHFR. Since L28R mutation is observed in almost 80 percent of all morbidostat trajectories and is synergistically interacting with several mutations, an L28R specific inhibitor will substantially impede evolution of trimethoprim resistance.

## Materials & Methods

### Growth Rate Measurements

All *DHFR* mutant strains were constructed in MG1655 attTn7::pRNA1-tdCherry (gift from Johan Paulsson). Detailed procedures for making mutant strains can be found [15]. Bacterial cultures were grown at 30 °C in M9 minimal medium supplemented with 0.4% glucose (Fisher Scientific B152-1), 0.2% amicase (MP Biomedicals 104778), 2mM MgSO_4_ (Fisher Scientific M63-500) and 100μM of CaCl_2_ (Fisher Scientific S25222A). Overnight grown cultures normalized to OD:0.001. Plates were incubated in 30°C with continuous shaking in Liconic Shaking Incubator and growth is measured with Tecan Plate Reader Infinite M200. Background optical density levels (OD∼0.04) are substracted from all wells. Growth rates are calculated by making an exponential fit to growth curve when bacterial growth is in its’ exponential phase.

### Intracellular DHFR abundance Measurements

*E. coli NDL47* cells were grown overnight, and final OD600 was adjusted to unity. These cells were then diluted by 10^4^-fold in 5 mL of M9 minimal media (supplemented with 0.4% glucose and 0.2% amicase) and grown for 6 h at 37°C (220 rpm) Cells were then washed three times with cold PBS buffer (pH 7.4), and bacterial pellets were lysed in 1X Laemmli sample buffer (5 mL/O.D.). Equivalent amounts of the cell lysates (10 μL of the above sample) from each set were electrophoresed in a 4%–15% precast polyacrylamide gel (561081; BIO-RAD), and western blotting was performed following standard procedures. DHFR antibodies are kindly provided by Kimberly Reynolds. IR-labeled secondary antibodies (IRDye 800CW (926–32213) and IRDye 680RD (925–68072); Li-COR) were used for detection. DHFR protein amount was quantified using an ODYSSEY infrared imaging system (LI-COR).

### Steady state Kinetic measurements

Reactants of DHFR reaction (DHF (Sigma-Aldrich D7006) and NADPH (Sigma-Aldrich N7505)) has absorbance at 340nm which the products (THF and NADP^+^) do not absorb light. Using LAMBDA 650 UV/Vis Spectrophotometer we measured reaction progression with 1sec resolution with two cells. First cuvette is sample cuvette containing the reaction components (DHFR, DHF and NADPH) and the second is reference cell contains only NADPH and DHFR in it. Biochemical measurements were done at 25°C in MTEN buffer (pH ∼7) which includes, 50mM MES hydrate (Sigma-Aldrich M8250), 25mM Tris-Base (Fisher Scientific B152-1), 25mM Ethanolamine Hydrochloride (Sigma-Aldrich E6133), 100mM NaCl (Fisher Scientific S271-3) and 5mM DTT (Fisher Scientific BP172-25) which is added fresh before starting the reaction. MTEN solution containing DHFR protein and 200 μM NADPH is prepared and 12.5μM DHF and 200μM NADPH solution is added preceding the data collection. Data collection is stopped when all the DHF is consumed which happens when the curve reach a plateau down below zero. Data analysis is done as explained in the main text (Figure 2A-B).

### Inhibition constant (*K*_*i*_) for TMP Determination

To calculate inhibition constants for TMP, we used initial rates of the reactions with saturating concentrations of DHF and NADPH with different TMP concentrations. These initial rates used to fit Michelis-Menten competitive inhibition formula to calculate *K*_*i*_ values (Figure 2C-D).

### Protein Overexpression and Purification

All combinations of six mutations of folA gene at five sites (I94L, W30R, W30G, L28R, A26T, P21L) are constructed by using Quick-Change Site-Directed Mutagenesis kit (Stratagene). 6XHis Tag is added on C-terminal of the protein sequence. Constructs are cloned into the expression plasmids (pET24a-KanR) for further protein purification. BL21 cells are transformed with pET24a-folA-6xHisTag were grown overnight in selective media (LB+Kan) and then diluted 100 times into TB media for further growth at 30°C. Protein overexpression induced when OD reached 0.6-0.8 using 250μM IPTG at 18°C with 220rpm shaking. Recombinant proteins are further purified using Ni-NTA columns (Qiagen) and dialyzed overnight using dialysis buffer containing 50mM Tris-Base, pH8.0, 0.5M NaCl, and 400mM Imidazole (Sigma Aldrich 792527).

### Epistasis Calculations and Linear Regression Model

A linear regression model is used to recover fitness of DHFR alleles by using epistatic interactions terms between DHFR mutations. The theory and algorithm we used to calculate epistatic terms and perform linear regression is described in detain by Poelwijk *et al.* [44].

### Molecular Dynamics Simulations

The NAMD package is used to model the dynamics of the protein–water systems [48]. Solvation is achieved *via* the VMD 1.9.1 program solvate plug-in version 1.2 [49]. The protein is soaked in a cubic solvent box such that there is at least a 10 Å layer of solvent in each direction from any atom of the protein to the edge of the box. The system is neutralized and 150 mM of ionic strength in all the simulations is maintained by adding a suitable number of K^+^ and Cl^−^ ions. The CharmM22 all-atom force field is used to model the protein and the TIP3P potential for water [41, 50]. We have adopted the force field parameters for 5-protonated 7,8-dihydrofolate and trimethoprim in two protonation states as reported in the literature [51]. Periodic boundary conditions are imposed on the simulation boxes that have 60 × 67 × 58 Å^3^ dimensions. Long range electrostatic interactions are calculated by the particle mesh Ewald method, [52] with a cutoff distance of 12 Å and a switching function at 10 Å. The RATTLE algorithm [53] is applied and a time step size of 2 fs in the Verlet algorithm is used. Temperature control is carried out by Langevin dynamics with a dampening coefficient of 5 ps^−1^. Pressure control is attained by a Langevin piston. All systems are first subjected to 10000 steps of energy minimization with the conjugate gradients algorithm. The resulting structures are then run in the NPT ensemble at 1 atm and 310 K until volumetric fluctuations are stabilized and the desired average pressure is maintained.

MD simulation of the ternary complex of the DHF bound systems are constructed based on the crystallographic structure with PDB code 1rx2 [37]. DHFR is complexed with folate and oxidized NADP (NADP^+^) in this native form. We protonate NADP and folate so that the former is in the reduced form (NADPH) and the latter is 5-protonated 7,8-dihydrofolate to model the stable state prior to the hydride transfer step.

In a separate set of MD simulations, we study the effect of trimethoprim binding in its unprotonated (TMP) or ground state (TMP^+^) on the DHFR conformation. Since there are no crystal structures of *E. coli* DHFR with trimethoprim, we have docked the inhibitor based on the coordination of equivalent residues of the trimethoprim binding region of *Staphylococcus Aureus* DHFR (PDB code: 2w9g) [38]. Details of trimethoprim binding site selection is provided in reference [41]. For MD simulations of the various mutants of DHF, TMP and TMP^+^ bound forms of DHFR, we mutated the WT structures *in silico via* BIOVIA Discovery Studio 4.0 package using build and edit protein tool [54]. For systems with multiple mutations, we substituted the native positions with the target mutations simultaneously. The solvation, ionization, minimization and equilibration were performed as described for the WT systems. All MD simulations are 210 ns long, with the first 10 ns discarded for equilibration. Simulations for the WT cases were repeated to confirm the reproducibility of the results.

The mutants studied are as follows: The single mutants I5F, M20I, P21L, A26T, D27E, L28R, W30G, W30R, I94L, R98P and F153S; all double mutant combinations of the A26T, L28R, I94L sets; the A26T-L28R-I94L triplet; the A26T-L28R-W30R-I94L and the A26T-L28R-W30G-I94L quadruplet. Also, to test the effect of the P21L mutation, we have studied the double mutant combinations of P21L with each of A26T, L28R, I94L as well as the P21L-A26T-L28R, P21L-A26T-I94L and P21L-L28R-I94L triplet, P21L-A26T-L28R-I94L quadruplet; and the P21L-A26T-L28R-W30R-I94L and the P21L-A26T-L28R-W30G-I94L quintets. Thus, we have carried out 210 ns long simulations of 26 sets of mutants, with DHF, TMP and TMP^+^ bound, leading to simulations exceeding 17.6 μs, including the WT sets.

We use the approach in reference [41] to confirm the native form of trimethoprim in the DHFR bound state, by monitoring the distribution of the native hydrogen bonds in the binding pockets. In all the sets, TMP^+^ remains tightly bound while TMP flips in and out of the binding pocket throughout the simulation. We thus accept the protonated form of trimethoprim to be the native form in all the systems; note that this is not the case for D27N and D27S mutants, as discussed at length in reference [41].

### Simulations of Protein Evolution and Visualization

Protein evolution simulations works on a DHFR mutational lattice (proteins as nodes and single mutation acquisition, conversion or reversion as lines). Simulations starts from WT in no trimethoprim condition. Trimethoprim concentration gradually increases and at each drug concentration fitness landscape of DHFR lattice is calculated. When drug concentration hits a value where enzyme activity is lower than threshold activity (50,10,1,0.1% of WT enzyme activity at [TMP] = 0 nM) a random mutational step is taken (a mutation acquisition, conversion or reversion). If the new mutant has lower activity than threshold, the simulation stops, otherwise the new mutation is fixed, and drug concentration starts increasing again till new mutants’ activity drops down to the threshold level (Figure 9B). Simulations are repeated for a million times to sample all possible unique trajectories. Python scripts to run the simulation is added to supplementary files. Visualization of the simulations is done by VPython, an open source software package for interactive 3D graphics [55].

## Acknowledgements

We would like to thank Roy Kishony, Adam Palmer, Shimon Bershtein, Adrian Veres, and Seungsoo Kim for their help.

---

**Figure S1: Comparison of predicted *V*_*0*_ and *K*_*i*_ values using Bliss additivity with experimentally measured values.** Panels on left side shows x-axis values predicted with a Bliss Independence model using single mutant data. Panels on the right shows x-axis values predicted with a Bliss Independence model using the (geometric) mean effects of single mutants.

**Figure S2: RMSF of inhibitor bound DHFR obtained from MD trajectories is increased as new mutations are added to the protein. A)** Maximum fluctuations in a residue are less than 3 Å in the WT protein (two replica simulations); the L28R single mutation and the L28R-I94L double mutation display their largest motions in the same regions as the WT, albeit with larger amplitude; with the addition of more mutations, largest amplitude motions are increased in size along with the introduction of additional mobile regions. **B)** The maximum RMSF values calculated for all the systems studied via MD are well correlated with experimentally measured log(*K*_*i*_) values for multiple mutant cases (filled circles; *r* = 0.71, *p* < 0.01) while they are uncorrelated for single mutants (*r* = 0.04, *p* > 0.9); overall correlations have *r* = 0.60; *p* < 0.01.

**Figure S3: Simulations are repeated for different threshold values (%*V*_*0*_ of WT as threshold) showing the number and length of evolutionary trajectories that reach to fitness peaks drastically increase if minimum fitness thresholds are assumed to be lower.**

**Table S1: WGS results of last days of morbidostat experiments show coding region mutations mostly occurred on folA gene (encodes DHFR).**

**Table S2: Mean measured *K*_*m*_, *k*_*cat*_, and *K*_*i*_ values of single mutants are shown in the table with standard error of the mean.** Additional sheets show measured replicates separately for each parameter (*K*_*m*_, *k*_*cat*_ and *K*_*i*_).

**Table S3: Summary of all measured *K*_*m*_, *k*_*cat*_, and *K*_*i*_ values for combination dataset.** This excel file also shows the Bliss Additivity calculations of *K*_*m*_, *k*_*cat*_, and *K*_*i*_ for both models with Single Mutant data and Mean Effects of Single Mutants.

**Table S4: Effects of addition of a single mutant in all backgrounds are shown as a table.** This data is plotted in Figure 6B.

**Table S5: Single mutants are appeared at different times in the morbidostat experiment.** Table shows the number of times each single mutant is appeared as the first coding sequence mutation. In the morbidostat experiment in almost all the cultures end with a single genotype dominating the culture. Second data column in this table shows the number of times a mutant is appeared in the last day of the evolution experiment.

**Table S6: Probability of seeing a mutant in the simulations are put in this table.** Each column shows different threshold (%*V*_*0*_ of WT as threshold) and whether the promoter mutation is allowed in the simulation.

